# Susceptibility of Simuliidae larvae (Diptera) to a new solid formulation based on *Bacillus thuringiensis* var. *israelensis* (Bti), and bioecological aspects of the breeding sites in Cascavel - Paraná, Brazil

**DOI:** 10.1101/2023.05.23.541956

**Authors:** Paula Costa Lis, Luis Francisco Angeli Alves, João Antônio Cyrino Zequi, Priscilla de Freitas Cardoso, Gislayne Trindade Vilas Boas, Francisco A. Marques, Renan R. Schorr, Itamar Francisco Andreazza, Óscar Sánchez Molina

## Abstract

Due to the hematophagous habits of females, some species of black fly (Diptera, Simuliidae) are known to be responsible for economic losses and can cause significant damage to the health of human and livestock populations. To control populations of these insects, several bioinsecticides based on the bacterium *Bacillus thuringiensis* var. *israelensis* (Bti) are widely used. The objective of the current study was to evaluate the susceptibility of black fly larvae to a new Bti solid effervescent formulation, comparing it with a commercial standard formulation, Vectobac® liquid formulation, under laboratory conditions. The study was carried out in the municipality of Cascavel, Paraná, Brazil. Eighteen hours after application of the formulations, larval mortality was evaluated. The control group did not exceed 20% mortality, for concentrations of 50 and 60 mg/L, the mean mortality rates were 50.6% and 64.2%, respectively, and neither differed significantly from the Vectobac®. The stream sampled showed external fecal contamination during the eight weeks of monitoring and water quality parameters that could interfere with the efficiency of the control with Bti at the site. The following species of black flies were collected and identified in the two watersheds sampled: *Simulium pertinax, S. subpallidum, S. nigrimanum, S. rubrithorax, S. perflavum*, and *S. inaequale*. Bioecological aspects of the breeding site were addressed and presented here, this being the first report of black fly species for the municipality of Cascavel. The potential of the solid effervescent formulation was proven under laboratory conditions and the bioecological evaluations demonstrated the relevance of environmental diagnosis and monitoring in the optimization of control protocols for Simuliidae.

## 1. INTRODUCTION

Black flies (Diptera: Simuliidae) are considered cosmopolitan (Duknić et al., 2019) and the immature stages have several morphological and physiological adaptations for development in lotic ecosystems (Adler and Mccredie, 2019). The larvae are key organisms in lotic ecosystems due to their ability to process dissolved organic matter, making it more readily available in the trophic chain (Malmqvist et al., 2004). On the other hand, the highly anthropophilic habits of some species can trigger irritative and allergic conditions, which can be associated with a low degree of well-being and economic losses in human activities, such as tourism and poultry, by affecting the quality of life and productivity of rural workers or in peri-urban areas associated with streams (Sá and Maia-Herzog, 2003). In addition, some species are vectors of several human and animal diseases (Crosskey, 1990). In Brazil, these insects are mainly related to the transmission of onchocerciasis and mansonellosis in humans (Shelley et al., 2010).

Moreover, in general, high densities of black fly populations are considered to be a consequence of deficiencies in environmental integrity, such as loss of biodiversity, reduction in natural predators, and high levels of organic matter present in anthropized and polluted streams. This excess of organic load in lotic environments may be a predisposing factor for the reduction in black fly diversity, which can also increase densities of singular species, for example the anthropophilic *Simulium pertinax* Kollar, 1832 (Strieder et al., 2006).

Faced with the problem of using chemical insecticides to control black fly populations, integrated management protocols containing *Bacillus thuringiensis* var. *israelensis* Berliner, 1911 (Bti) are currently used. This bacterium, which can be found naturally in the soil of different regions of the world, has a parasporal body (crystal) composed of proteins that, when ingested by the black fly larvae, has the potential to destroy midgut epithelial cells, leading to sepsis and death (Bravo et al., 2007). The Bti-based bioinsecticides available in Brazil are imported and demand high investment from the consumer. Therefore, the development of national formulas, with a low acquisition cost when compared to imported products, would optimize their use in integrated management programs for black fly control (Angelo et al., 2010).

On the other hand, good control results also require actions developed considering technical criteria, including as a minimum, knowledge of the environmental conditions of the treated site. The application intervals must be respected, as it is known that smaller black fly larvae instars predominate in sections of streams treated with Bti when compared to untreated streams, meaning that there is constant recolonization of treated streams (Docile et al., 2021). The indiscriminate establishment of protocols can lead to the risk of extinction of Simuliidae species at treated points (Cheke et al., 2008), while the lack of criteria leads to product waste (Rivers-Moore et al., 2008).

Local production and obtaining national formulas, with a low acquisition cost when compared to imported products, would optimize their use in integrated management programs in the control of black flies (Bhumiratana, 1990; Ampofo, 1995). The use of solid presentations for the control of black flies, presented in this work, is innovative, and aims to facilitate the application process, especially in areas of difficult access, envisioning the possibility of obtaining an effervescent product that can be applied directly to the river by the affected population itself (Becker, 2000). Effervescent tablets based on Bti (Culinex®), have been used since 1992 with efficacy against *Culex pipiens* Linnaeus, 1758. Thus, the current work presents the evaluation of a new solid formulation for the control of black fly. In addition, we also describe the bioecological aspects of breeding sites in streams of Cascavel, Paraná, Brazil and the species of Simulium found in the evaluated basins. Although there are records of species of *Simulium* for the state of Paraná (Lozovei, 2004), in the municipality of Cascavel there are still no records of the presence of the species described in the literature.

Therefore, the present study aimed to evaluate the susceptibility of the larvae of *Simulium* to an effervescent Bti formulation, under laboratory conditions, and correlate it to the bioecological aspects of the breeding sites.

## 2. MATERIAL AND METHODS

### 2.1 Larvae and water river collection

Larvae collections were carried out with authorization from the Chico Mendes Institute for Biodiversity Conservation – ICMBio – registered under number 73993-1 in the Biodiversity Authorization and Information System – SISBIO, Ministry of the Environment – MMA. The insects were maintained in the laboratory in a controlled room (25±2°C), with permanent aeration maintained by two-outlet air compressors (Boyu ®, SC – 7500 - 2×3L/min, 127 V).

### 2.2 Bioinsecticide - Bacteria

To produce the bioinsecticide, the *Bacillus thuringiensis* subsp. *israelensis* (Bti) strain BR101 was used, from the Entomopathogenic Bacteria Bank of the Laboratory of Bacterial Genetics and Taxonomy of the State University of Londrina - UEL, Department of Animal and Plant Biology, of the Center of Biological Sciences (CCB) – Londrina PR.

Cultures of *Bacillus thuringiensis* strain BR101 were performed in a bioreactor containing 5L of NYSM medium (Nutrient Yeast Extract Salt Medium) and maintained at 30° C ± 2,300 rpm (rotations per minute), pH 7.0 (hydrogenonic potential), with constant air supply, for about 40 hours, until sporulation. Subsequently, the fermentation broth was centrifuged, the supernatant discarded and the final suspension was lyophilized.

#### 2.2.1 Formulations

The effervescent solid dosage forms were prepared at the Laboratory of Chemical Ecology and Synthesis of Natural Products and Pharmaceutical Technological Laboratory of the Federal University of Paraná (UFPR), in Curitiba, Paraná, Brazil. They were prepared by direct compression with a manual hydraulic press (Maxx press, Essence dental-SP), using a flat and circular punch, with a diameter of 11 mm. The compression force was standardized at 500 kgf for 20 seconds until the desired pressure was reached, and then maintained for another 10 seconds with 500 kgf.

Seven formulations were prepared (Table 1), keeping constant the presence of the effervescent mixture (citric acid, sodium bicarbonate, and sodium carbonate) as disintegrant, sucrose as soluble diluent, and powdered sodium lauryl sulfate as wetting/disintegrant. Corn starch was used as a diluent in varying concentrations to adjust the final mass of the tablet, established at 250 mg. The lyophilized solid containing the Bti protein crystals was used in increasing amounts (Table 1). The materials were weighed and hand mixed with low friction (plastic bag). After this process, compression was carried out, weighing the mass individually, using 250 mg of the corresponding mixture for each tablet produced.

**Table 1.** Composition and quantity of ingredients in solid effervescent tablets (mg) based on Bacillus *thuringiensis* var. *israelensis*.

| Ingredients | Formulations |  |  |  |  |  |  |
| --- | --- | --- | --- | --- | --- | --- | --- |
|  | 1 | 2 | 3 | 4 | 5 | 6 | 7 |
| Effervescent mixture | 105.55 | 105.55 | 105.55 | 105.55 | 105.55 | 105.55 | 105.55 |
| Starch | 42.45 | 39.45 | 34.45 | 29.45 | 24.45 | 19.45 | 14.45 |
| Sucrose | 90.00 | 90.00 | 90.00 | 90.00 | 90.00 | 90.00 | 90.00 |
| *Lyophilized solid | 2.00 | 5.00 | 10.00 | 15.00 | 20.00 | 25.00 | 30.00 |
| Sodium Lauryl sulfate | 10.00 | 10.00 | 10.00 | 10.00 | 10.00 | 10.00 | 10.00 |
| TOTAL (mg) | 250.00 | 250.00 | 250.00 | 250.00 | 250.00 | 250.00 | 250.00 |

\*Solid containing proteins with larvicidal activity

#### 2.2.2 Effervescent mix

The effervescent mixture was prepared at the time of final mixing (Prista et al, 1967); for every 10 parts of citric acid, 12 parts of sodium bicarbonate and 7.55 parts of sodium carbonate were used.

#### 2.2.3 Standard Formulation

The commercial product Vectobac® 12 AS (liquid formulation) with 1,200 UTI/mg (Lot 301-886-N930 – Expiration date: 05/2021), was used as a comparison standard for the test formulations. The suggested concentration for the product is from 0.5 to 25 ppm/L for 1 minute of exposure. However, in the tests, the amount of 0.2 mL/0.5L (400 ppm) was stipulated as standard. This concentration was defined above that recommended by the manufacturer, considering the ease of application, and thus avoiding the need for serial dilutions. In addition, the product was also used with the objective of validating the applied methodology, by evaluating the response to the presence or absence of mortality due to Bti, with no intention of testing the potency of the commercial product.

### 2.3 Bioassays

The bioassays were carried out based on Araújo and Coutinho et al. (2005) with modifications. Larvae were placed in plastic containers containing water collected directly from the environment where they were collected. Active searches for larvae, to perform the bioassay for the evaluation of larvicides, were carried out in seven streams in the urban and rural areas of the city of Cascavel, Paraná State, Brazil.

Among all the sample points studied, the Santa Rosa stream was selected to carry out the bioassays, as it has the highest abundance of blackflies larvae and was not being treated with bioinsecticides or other products to control them. This stream is in the Sede Alvorada district, a rural area 23 km from the municipality of Cascavel. It is 7 km long and is a tributary of the São Francisco Verdadeiro river (24°49’15.9” S and 53°37’51.2” W). Only larvae collected on the same day as the test were used, thus preventing the laboratory time from interfering more significantly with their mortality.

In the laboratory, the larvae were evaluated for motility and viability, avoiding those with dark gill histoblasts, as this is a characteristic of the last stage of development, which would present a risk of the larva pupating during the experiment. The material was transferred to plastic containers filled with 400 ml of water from the collection point, individually and uninterruptedly oxygenated with an aerator compressor. After an acclimatization period of 3 hours, the larvae that had not been set were discarded and replaced on-site, ensuring that only healthy larvae were evaluated (Lacey, 1997). Because of the source of the water, no external nutritional compounds were applied. The water temperature during collection, transport, and the experiment was maintained at between 20 and 25°C.

For the application of the solid formulations, the tablets were diluted in 100 mL of water from the river itself. The mixture was applied to the respective recipients and 1 minute was allowed for the larvae to ingest the Bti. Subsequently, the same procedures described for the formulated commercial liquid were adopted.

For each of the test products and for the control, three cups were prepared, each considered a repetition. The mortality of 20% of the control group was accepted as a cut-off point, since the larvae are sensitive to the process of collection, handling, transport, and the laboratory environment (Lacey, 1997).

### 2.4 Data analysis

All bioassays were conducted in a completely randomized design. Larvae mortality data were submitted to analysis of variance and the Shapiro-Wilk (p<0.05) residual normality test. The homogeneity of variance was then evaluated using the Bartlett test (p<0.05) and the means were compared using the Tukey test (p<0.05).

### 2.5 Taxonomic identification

As taxonomic identification is a premise for carrying out control programs of the species that inhabit the streams targeted by the actions, part of the material collected for the bioassay was sent to the Laboratory of Simulids and Onchocerciasis of the Instituto Oswaldo Cruz (LSO/IOC), for taxonomic identification.

### 2.6 Characterization of the bioecological parameters of the selected collection point

To assess the water quality parameters at the point of the bioassay, data collections were carried out to analyze the physical-chemical and microbiological characteristics of the water, weekly from 03/10/2021 to 04/27/2021. These data were analyzed at the Foundation for Scientific and Technological Development (FUNDETEC - Cascavel).

The physicochemical parameters analyzed were the hydrogenic potential (pH), electrical conductivity, turbidity, total solids, nitrite, nitrate, and phosphate. The microbiological analysis investigated the presence of total coliforms and *Escherichia coli* (Escherich, 1885). Water temperature, air temperature, and relative air humidity were observed at the time of water collection. To verify the integrity of the stretch, the Protocol for the Rapid Assessment of Habitat Diversity in stretches of hydrographic basins (PAR) by Callisto et al. (2002), and adapted from Hannaford et al., (1997), was applied on site. This protocol aims to assess the conditions observed in the basin stretch, anthropic influence, habitat conditions, and level of conservation, assigning a score for each characteristic observed in the location. The sum of points represents whether the stretch is impacted (0 to 40 points), altered (41 to 60 points), or natural (above 60 points).

## 3. RESULTS

### 3.1. Bioassay

The concentration of 50 mg/L presented a mean mortality of 50.6%, which did not differ significantly from the commercial standard formulation, or from the concentration of 40 mg/L. For the concentration of 60 mg/L, significant mortality was observed, with an average of 64.2%, not significantly differing from the standard formulation. The commercial standard formula presented a mean mortality of 75.4% after the evaluation time of 18 hours (p<0.05) (**Table 2**). Concentrations of 4 mg/L, 5 mg/L, and 10 mg/L did not show significant mortality.

**Table 2.** Average mortality of Simuliidae larvae, submitted to the treatment of solid larvicides based on

| Treatment | Average mortality (%) |
| --- | --- |
| Standard formula | 75.4 a |
| 60 mg/L | 64.2 a |
| 50 mg/L | 50.6 ab |
| 40 mg/L | 34.3 bc |
| 30 mg/L | 19.6 c |
| Control | 11.3 c |

Means ( ± SEM) followed by the same letter do not differ statistically from each other, using the Tukey test ( $p < 0.05$ )

### 3.2. Water parameters and physical integrity of the collection site

Weekly oscillations were observed in the parameters evaluated, mainly in the amount of Total Solids, with a minimum value of 72.1 and maximum of 423.9 ppm (CV% = 70.8) and in the parameters of Nitrate, from 0.9 to 1.6 ppm (CV% = 16.8) (**Table 3**). The presence of total coliforms and *Escherichia coli* was observed in all collections performed.

**Table 3.** Physicochemical conditions of the water of Santa Rosa Stream, Paraná Basin 3, Cascavel, Paraná, Brazil. Monitored weekly from 03/10/2021 to 04/28/2021.

| Variables | Mean value | CV (%) |
| --- | --- | --- |
| pH | 7.2 ± 0.5 | 6.5 |
| Conductivity (µS/cm) | 17.3 ± 0.9 | 5.1 |
| Turbidity (NTU) | 17.4 ± 2.2 | 12.9 |
| Total Solids (ppm) | 174.1± 123.2 | 70.8 |
| Nitrite (ppm) | < 0.01 | 0 |
| Nitrate (ppm) | 1.4 ± 0.2 | 16.8 |
| Phosphate (ppm) | 8.4 ± 0.5 | 6 |
| Water temperature (°C) | 19.8 ± 1.3 | 6.8 |
| Temperature (°C) | 20.8 ± 1.3 | 6.2 |
| Relative humidity % | 70.6 ± 7.4 | 10.5 |

\* Means (±EPM); CV: Variability coefficient;; pH: potential of hydrogen; µS/cm: micro-Siemens per centimeter; NTU: Nephelometric Turbidity Unit; ppm: parts per million.

In the investigated stretch of the Santa Rosa stream, the Protocol for the Rapid Assessment of Habitat Diversity in stretches of hydrographic basins (Callisto et al., 2002) indicated that the stretch can be characterized as a natural environment.

### 3.3. Taxonomic identification

Altogether, 22 exuviae, 55 pupae, ten adults, 42 mature larvae, and 299 immature larvae were collected and examined. Six species belonging to the genus *Simulium* Latreille, 1802 were identified and distributed in five subgenera: *Simulium* (*Chirostilbia*) Enderlein, 1921 (two spp.), *S*. (*Hemicnetha*) Enderlein, 1934 (one sp.), *S*. (*Psaroniocompsa*) Enderlein, 1934 (one sp.) *S*. (*Psilopelmia*) Enderlein, 1934 (one sp.), and *S*. (*Trichodagmia*) Enderlein, 1934 (one sp.) (**Table 4**).

**Table 4.**
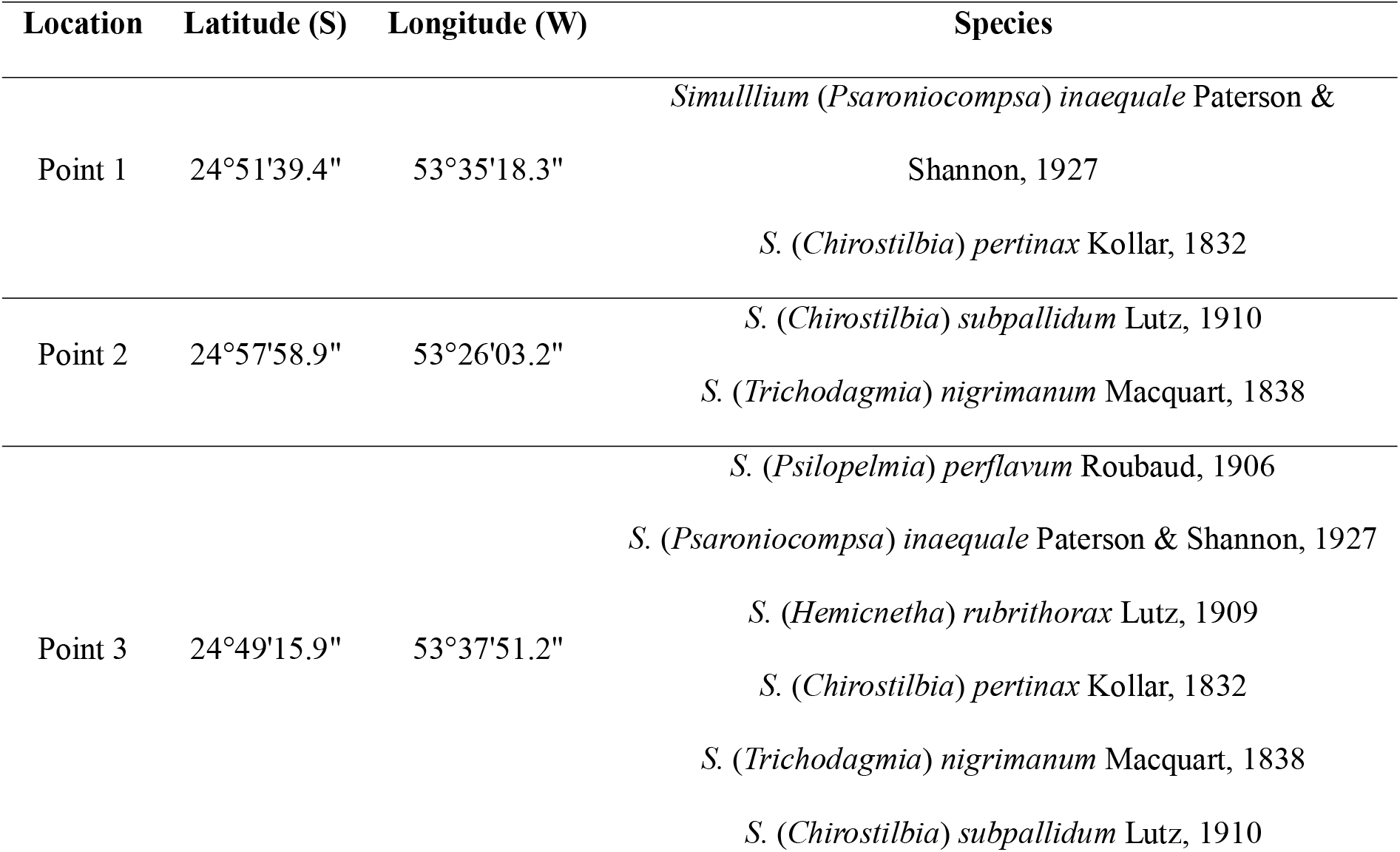
Black fly species found at sampled points in Cascavel, PR, Brazil. 2019 -2020.

At points 1 and 2, only two species were found, whereas at Point 3 the six species identified in this study were present, this being consequently the point of greatest diversity (**Table 4**). However, this may have been due to the higher number of samples collected at this location.

## 4. DISCUSSION

### 4.1. Larvicidal activity of bioinsecticides

The exposure time of 1 minute was used, associated with high concentrations of Bti. However total larval mortality was not observed in any of the treatments, which can be explained by the reduced exposure time.

Histological alterations in midgut epithelium were observed in *S. pertinax* larvae after 1 hour of Bti exposure (6 mg/L), in addition to total larval mortality (Cavados et al. 2004), however this exposure time is much longer than usual for field applications.

### 4.2. Water parameters and physical integrity of the collection site

Regarding water parameters, in an experiment conducted by Wilson et al. (2005) the best mortality results of a Bti-based product against *S. damnosum* Theobald, 1903 larvae were observed as the temperature increased. According to the authors, temperature influences metabolism and, consequently, greater ingestion of toxins by larvae. Atwood et al. (1992), in an experiment conducted with larvae of *Cnephia pecuarum* (Riley 1887), observed a reduction in larval mortality in response to Bti when the water temperature decreased, recommending that the control be carried out before the temperature drops below 9 °C. Therefore, the same recommendation could be applied to the Santa Rosa stream, especially in the winter season, with the measurement of this parameter and application of Bti only if the water temperature is above 9°C.

Wilson et al. (2005) found that the high conductivity of the water provided better larvicidal performance of the product, obtaining the best result in the conductivity of 323 mS (millisiemens), much higher than that observed in the Santa Rosa stream, with an average conductivity of 16.2 to 19.1 μS/cm. According to the authors, most streams do not have such high conductivities under natural conditions, and this parameter should not be a decisive factor in the use of the product.

For the turbidity parameter, Wilson et al., (2005) observed better responses to Bti when the water turbidity was below 4.2 Nephelometric Turbidity Units (NTU), and recommended that, for the operational use of the product, turbid water should be avoided. Therefore, if a simulid control protocol was instituted in the evaluated stream, it would be important to carry out monitoring of the water quality and rainfall, comparing the information with the larvicidal efficiency of the product, since the water parameters showed oscillations due to rainfall and discharges of organic matter.

In the Santa Rosa stream, the mean variation in nitrate was 0.9 to 1.6 ppm and *S. pertinax* was identified, which is similar to the results of a study carried out in an impacted basin in southern Brazil, where a correlation was found between the occurrence of *S. pertinax* and nitrate concentrations from 1.0 mg/L to 1.75 mg/L, related to streams with a greater organic pollution and environmental degradation in poultry and swine production sites (Strieder., 2006).

The averages of the variables pH, turbidity, total solids, nitrite, and nitrate in the Santa Rosa Stream water were framed in CONAMA Resolution No. 357, of March 17, 2005, as values observed for Class 1 of freshwater quality standards (which is the best classification standard and can be used for human consumption after disinfection). However, the microbiological analysis indicated the presence of total coliforms and the bioindicator *Escherichia coli*, so that the Santa Rosa stream presented external fecal contamination during the eight weeks of monitoring, which could be related to the cattle and other animals being raised on nearby properties. It is also believed that external contamination may be the source of organic matter for the maintenance of immature breeding sites in the stream.

Although the Protocol for the Rapid Assessment of the Diversity of Habitats in stretches of hydrographic basins (PAR) presented 66 points, which is configured as a “natural” stretch, several alterations were diagnosed in the stream, including the presence of contamination by thermotolerant coliforms, erosion, and deforestation. These alterations, if they persist, may change the score of the section in the future, considering that it is already very close to being classified as an altered environment (60 points). Therefore, the association between loss of water quality and the presence of breeding sites could guide the community to get involved and take responsibility for reducing environmental conditions that generate imbalance, arising from the activities carried out in the localities, as well as generating information that will be useful in the formulation of public policies, mainly environmental and health.

### 4.3. Considerations on Taxonomic Identification

Among the identified species, some stand out for their medical, veterinary, or socioeconomic interest. *Simulium nigrimanum* Macquart, 1838 is anthropophilic, being a vector of Onchocerciasis in the Brazilian Minaçu focus. It has also been associated with the development of the autoimmune human disease known as *pemphigus foliaceous*, characterized by the appearance of extensive bullous lesions and cutaneous erosions (Eaton et al. 1998; Shelley et al., 2010).

The females of *Simulium inaequale* Paterson & Shannon, 1927 are highly anthropophilic, however, they have also been recorded biting mules (Shelley et al., 2010). *Simulium pertinax* is the most common anthropophilic species in southern Brazil and has also been recorded biting horses in Brazil and cows in Bolivia (Shelley et al., 2010). This species can be found at high densities in anthropic streams, being the main target in several simulid control programs. The presence of this species at Point 3 may be associated with the nitrate variations found (Table 1), which agrees with the result of Strieder et al., (2006), who found a significant association of nitrate concentration with the presence of *S. pertinax*.

Furthermore, females of *S. perflavum* Roubaud, 1906, *S. subpallidum* Lutz, 1910, and *S. rubrithorax* Lutz, 1909 are zoophilic, the latter having been recorded feeding on horses and probably cattle (Coscarón et al., 2007, Shelley et al., 2010). The feeding habit of *S. rubrithorax* is compatible with the place where it was collected, Point 3, as the presence of production animals, such as cattle, was observed on nearby properties.

Although simulids are widely known for their nuisance potential, their larvae are key organisms in aquatic ecosystems as an important food source for vertebrate and invertebrate organisms, and, due to their ability to process dissolved organic matter, making it more readily available in the food chain (Malmqvist *et al*., 2004). In addition, numerous species that do not interfere with humans or their activities are also affected by control methods, reinforcing the need for previous knowledge of the biodiversity of species existing in target locations of control, in order to subsidize sustainable and efficient actions (Malmqvist et al., 2004).

This is the first record of Simuliidae for the city of Cascavel and considering that the collections were carried out in only three fragments of lotic environments, and only two of the three hydrographic basins, it is believed that the diversity of species in the urban and rural areas of the city may be greater.

## 5. Conclusion

The innovative use of a solid Bti compound for black fly control presented in this work, aims to facilitate the product application process, especially in areas of difficult access, envisioning the possibility of obtaining an effervescent product that can be applied directly to the water course without the need for prior dilution. The data obtained indicate the larvicidal potential of this new effervescent formulation, with proven efficacy in the laboratory.

The present work also demonstrates that in the study area there is biodiversity of black fly fauna, of human and veterinary health importance, associated with urban and rural streams. The bioecological assessments reinforce the relevance of environmental diagnosis and monitoring in the optimization of Simuliidae control protocols.

## References

ABNT - Associação Brasileira De Normas Técnicas., 1987. NBR 9898 - Preservação e técnicas de amostragem de efluentes líquidos e corpos receptores. Rio de Janeiro.

Adler, P.H., Mccreadie, J.W., 2019. Black flies (Simuliidae), in: Mullen, G.R., Durden, L.A. (Eds.), Medical and Veterinary Entomology, New York, Academic Press, pp. 237–259.

Aoki, V., Rivitti, E.A., Ito, L.M., Hans-Filho, G., Diaz, L. A., 2005. Historical profile of the immunopathogenesis of endemic pemphigus foliaceus (fogo selvagem). An. Bras. Dermatol. 80, 287–292. https://doi.org/10.1590/S0365-05962005000300010.

Araújo-Coutinho, C.J.P.C., Figueiró, R., Viviani, A.P., Nascimento, E.S., Cavados, C.F.G., 2005. A bioassay method for black flies (Diptera: Simuliidae) using larvicides. Neotrop. Entomol. 34, 511–513. https://doi.org/10.1590/S1519-566X2005000300022.

Atwood, D.W., 1992. Efficacy of Bacillus thuringiensis var. israelensis against larvae of the southern buffalo gnat, Cnephia pecuarum (Diptera: Simuliidae), and the influence of water temperature. J. Am. Mosq. Control Assoc. 8, 126–30. https://pubmed.ncbi.nlm.nih.gov/1431853/.

Bocaleti, A.S., 2019. Novos formulados à base de Bacillus thuringiensis subesp. israelensis para controle de Culicidae (Diptera). 2019. 65 f. Dissertação (Mestrado em Ciências Biológicas) – Universidade Estadual de Londrina, Londrina – Paraná.

Bravo, A., Gill, S. S., Soberón, M., 2007. Mode of action of Bacillus thuringiensis Cry and Cyt toxins and their potential for insect control. Toxicon 49, 423–35. https://doi.org/10.1016/j.toxicon.2006.11.022.

Brasil. Conselho Nacional do Meio Ambiente – CONAMA, 2005. Resolução CONAMA N° 357.17/03/2005.Brasília,DFhttp://pnqa.ana.gov.br/Publicacao/RESOLUCAO_CONAMA_n_357.pdf

Callisto, M. Ferreira, W. R., Moreno, P. Goulart, M. Petrucio, M., 2002. Aplicação de um protocolo de avaliação rápida da diversidade de hábitats em atividades de ensino e pesquisa (MG-RJ). Acta Limnol. Bras. 14, 91–98. http://jbb.ibict.br//handle/1/708.

Cavados, C.F.G., Majerowicz, S., Chaves, J. Q., Araújo-Coutinho, C. J. P. C., Rabinovitch, L., 2004. Histopathological and ultrastructural effects of delta-endotoxins of Bacillus thuringiensis serovar israelensis in the midgut of Simulium pertinax larvae (Diptera, Simuliidae). Mem. Inst. Oswaldo Cruz 99, 493–498. https://doi.org/10.1590/S0074-02762004000500006.

CETESB - Companhia Ambiental do Estado de São Paulo, 2011. Guia Nacional de Coleta e Preservação de Amostras - Água, Sedimento, Comunidades Aquáticas e Efluentes Líquidos.

Coscarón, S., Coscarón-Arias, C.L., 2007. Neotropical Simuliidae (Diptera: Insecta), in Adis, J., Arias, J.R., Rueda-Delgado, G., Wantzen, K.M. (eds.) Aquatic biodiversity in Latin America, Pensoft, Sofia-Moscow, 685 pp.Biodiversidade aquática na América Latina, Pensoft, Sofia-Moscou.

Crosskey, R.W., 1990. The natural history of blackflies. Ltd. Baffins Lane. Chichester.Docile, T.N., Figueiro, R., Gil-Azevedo, L.H., Nessimian, J.L., 2015. Water pollution and distribution of the black fly (Diptera: Simuliidae) in the Atlantic Forest. Brazil. Int. J. Trop. Biol. 63, 683–693. https://www.scielo.sa.cr/scielo.php?script=sci_arttext&pid=S0034-77442015000300683.

Duknić, J., Jovanovic, V. M., Popvic, N., Zivic, I., Rakovic, M., Cerba, D., Paunovic, M., 2019. Phylogeography of Simulium ubgenus Wilhelmia (Diptera: Simuliidae) - Insights from Balkan Populations. J. Med. Entomol. 56, 967–978. https://doi.org/10.1093/jme/tjz034.

Eaton, D.P., Diaz, L. A., Hans-Filho, G., Santos, V. D., Aoki, V., Friedman, H., Rivitti, E. A., Sampaio, S. A., Gottlieb, M. S., Giudice, G. J., Lopez, A., Cupp, E. W., 1998. Comparison of Black Fly Species (Diptera: Simuliidae) on an Amerindian Reservation with a High Prevalence of Fogo Selvagem to Neighboring Disease-Free Sites in the State of Mato Grosso do Sul, Brazil. J. Med. Entomol. 35, 120–131. https://doi.org/10.1093/jmedent/35.2.120.

Lacey, L. A., 1997. Bacteria: Laboratory bioassay of bacteria against aquatic insects with emphasis on larvae of mosquitoes and black flies, in: Lacey, L.A. (Ed.), Manual of Techniques in Insect Pathology, Academic Press, Cambridge, pp.79–90.

Maia, A., Direito, I.C.N., Figueiró, R., 2014. Controle biológico de simulídeos (Diptera: Simuliidae): panorama e perspectivas. Cadernos UniFOA, 9, 89–104. https://doi.org/10.47385/cadunifoa.v9.n25.116.

Maia-Herzog, M., Shelley, A.J., Dias, A.P.A.L., Malaguti, R., 1984. Comparação entre Simulium Brachycladum e S. Rubrithorax, suas posições no subgênero Hemicnetha e notas sobre uma espécie próxima, S. Scutistriatum (Díptera: Simuliidae). Mem. Inst. Oswaldo Cruz, 79, 341–356. https://doi.org/10.1590/S0074-02761984000300008.

Malmqvist, B. Adler, P. H., Kuusela, K., Merritt, R. W., Wotton, R. S., 2004. Black flies in the boreal biome, key organisms in both terrestrial and aquatic environments: A review. Ecoscience, 11, 187–200. http://www.jstor.org/stable/42901602.

Rio Grande do Sul., 2018. Secretaria Estadual da Saúde. Centro Estadual de Vigilância em Saúde. Vigilância Ambiental de Simulídeos (Díptera, Simuliidae) no Rio Grande do Sul: Orientação para gestão nos municípios. Porto Alegre CEVS/RS. https://www.cevs.rs.gov.br/upload/arquivos/201910/18110908-2018-caderno-de-simulideos.pdf

Sá, M.R., Maia-Herzog, M., 2003. Overseas disease: comparative studies of onchocerciasis in Latin America and África. Hist. Cienc. Saude Manguinhos, 10, 251–258. https://doi.org/10.1590/S0104-59702003000100008

Shelley A.J., Hernández, L.M., Maia-Herzog M., Luna, DA,, Garritano, P.R., 2010. The Blackflies of Brazil (Diptera, Simuliidae), in: Adis, J., Arias, J., Golovatch, S S., Mantzev, K.M., Rueda-Delgado, G., Domínguez, E. (Eds.), Aquatic Biodiversity in Latin America (ABLA Series), Pensoft, Sofia-Moscosw.

Strieder, M.N., Santos, J. E., Vieira, E. M., 2006. Distribution, abundance and diversity of Simuliidae (Díptera) in an impacted watershed in southern Brazil. Rev. Bras. Entomol. 50, 119–124. https://doi.org/10.1590/S0085-56262006000100018.

Wilson, M.D., Akpabey, F. J., Osei-Atweneboana, M. Y., Boakye, D. A., Ocran, M., Kurtak, D. C., Cheke, R. A., Mensah, G. E., Birkhold, D., Cibulsky, R., 2005. Field and laboratory studies on water conditions affecting the potency of VectoBac® (Bacillus thuringiensis serotype H-14) against larvae of the blackfly, Simulium damnosum. Med. Vet. Entomol. 19: 404–412. https://doi.org/10.1111/j.1365-2915.2005.00591.x.

